# Decoding mTOR signalling heterogeneity in the tumour microenvironment using multiplexed imaging and graph convolutional networks

**DOI:** 10.1101/2023.12.30.573693

**Authors:** Razan Zuhair, Mark Eastwood, Megan Jones, Amy Cross, Joanna Hester, Fadi Issa, Fiona Ginty, Heba Sailem

**Affiliations:** School of Cancer and Pharmaceutical Sciences, Stamford St., Franklin Wilkins Building, King’s College London, London SE1 9NQ; Tissue Image Analytics Center, University of Warwick; Nuffield Department of Surgical Sciences, Level 6, John Radcliffe Hospital, University of Oxford, Oxford OX3 9DU; GE Research, Niskayuna, NY, USA; Department of Engineering Science, Old Road Campus Research Building, University of Oxford OX3 7DQ, UK

**Keywords:** Tumour Microenvironment, Multiplexed Imaging, mTOR signalling, graph neural networks, spatial biology

## Abstract

Evaluating the contribution of the tumour microenvironment (TME) in tumour progression has proven a complex challenge due to the intricate interactions within the TME. Multiplexed imaging is an emerging technology that allows concurrent assessment of multiple of these components simultaneously. Here we utilise a highly multiplexed dataset of 61 markers across 746 colorectal tumours to investigate how complex mTOR signalling in different tissue compartments influences patient prognosis. We found that the signalling of mTOR pathway can have heterogeneous activation patterns in tumour and immune compartments which correlate with patient prognosis. Using graph neural networks, we determined the most predictive features of mTOR activity in immune cells and identified relevant cellular subpopulations. We validated our observations using spatial transcriptomics data analysis in an independent patient cohort. Our work provides a framework for studying complex cell signalling and reveals important insights for developing mTOR-based therapies.

## Introduction

Recent advances in multiplex tissue imaging provide an unprecedented opportunity for studying complex cell signalling because it permits investigating multiple components in the tumour and tumour microenvironment (TME) simultaneously at a single-cell level^1–4^. For example, the identity of different cell types can be visualised in context of their spatial organisation, signalling state and morphologies. This capability allows profiling signalling activities in various interacting cell types including cancer, immune and stromal cells which enable a better understanding of how heterocellular cell signalling drives cancer progression and impacts patient outcome. Such understanding is crucial for designing effective targeted treatment strategies that account for the interaction between different TME components.

The mTOR pathway integrates nutritional and environmental cues to modulate cell proliferation, metastasis, metabolism. It has a multifaceted influence on the TME by inducing angiogenesis and immune chemotaxis thereby exerting a multifaceted influence^5–7^. Dysregulation of mTOR in fibroblasts can support tumour invasion and fibroblast activation^8,9^. Moreover, mTOR also plays a vital role in immune response and immune cell proliferation and differentiation including Tregs, CD8+ T-cells, macrophages, neutrophils and B-cells^10–15^. However, simultaneous analysis of mTOR activities in cancer cells and the surrounding immune microenvironment and how they drive cancer progression in different tumour microenvironments is underexplored. Moreover, how immune mTOR activity impacts cancer cell signalling and fate largely remains unanswered.

The activation of mTOR, leads to activation of ribosomal protein S6 and inhibition of eukaryotic translation initiation factor 4EBP1 through its phosphorylation. S6 phosphorylation has been associated with worse patient prognosis^16,17^. However, there have been conflicting results on the impact of the 4EBP1 phosphorylation on survival as it has been associated with both worse and better prognosis^17–19^. In this study, we hypothesised that the spatial effects of mTOR in activity in immune cells on cancer cells might have an effect on the overall impact of mTOR on patient prognosis.

Representing tumours as a network of interacting cells is an effective strategy for considering spatial protein activity and tumour-microenvironment interactions. Graph convolutional network (GCN) is particularly suited for learning from graph structured data as it takes a graph or a network as an input^20^. GCNs inherently capture the interacting nature of cells through message-passing procedure. This allows capturing several levels of influence, as at each GCN layer, cells update their state to reflect their own state but also their neighbours’ state. After each iteration, the states of each cell’s neighbours also incorporate the states of their own neighbours which enable capturing short- and long-range cellular communications. Another notable advantage of GCNs over traditional deep learning approaches is explainability as they allow for the identification of the contributions of cellular subpopulations to specific signaling states.

In this study, we utilised multiplexed imaging data of 746 colorectal tumours to profile mTOR activity. In addition to markers of various proteins in the mTOR pathway, we included markers related to key cancer pathways to determine interconnected signalling cascades. We performed compartmentalised image analysis of protein activities in epithelial, immune and stromal cells to study how mTOR activity in different cell types affects patient outcomes. We combined unsupervised machine learning with an explainable graph neural network to effectively interrogate quantitative single cell signalling data. We found that mTOR can play a differential role in cancer progression depending on the tumour microenvironment. Our results underscore the importance of considering complex protein signalling in the TME.

## Results

### Compartmentalised image analysis reveals differential mTOR association with patient prognosis

To investigate mTOR activity and associated signalling and TME phenotypes in colorectal cancer, we utilised highly multiplexed data where cyclic immunofluorescence was used to image the activity of 61 markers. 746 colorectal tumours with different stages and grades were imaged^1^ (Fig 1A-B and Methods). These markers encompassed various structural and TME markers, including epithelial markers (pan-cytokeratin (PCK), E-cadherin, EPCAM and beta-actin), extracellular matrix markers (fibronectin and collagen), stromal markers (smooth muscle actin (SMA) and Vimentin) and immune markers (CD3, CD8, CD20, CD68, and CD79). Additionally, these sections were stained for several markers of mTOR pathways including 4EBP1, p4EBP1, S6, pS6 and the upstream signalling pathways AKT, PTEN, p38, EGFR, MET and ERK. (Fig. 1B and Supplementary Table 1).

**Fig. 1.**
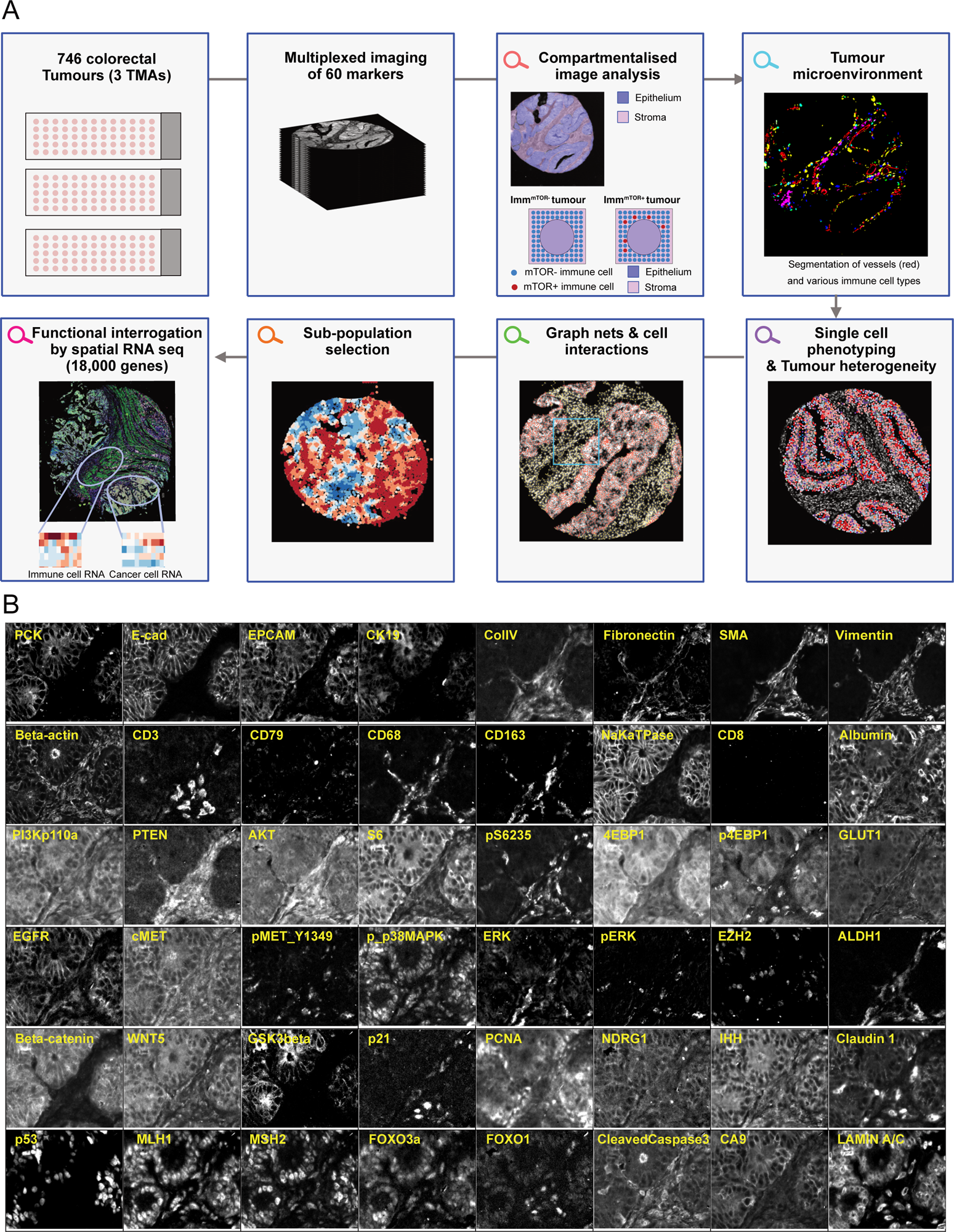
Multiplexed imaging allows multi-level investigation of complex cell signalling. A) Overview of the study with many of the indicated analyses can be performed concurrently. Immune mTOR status in tumours is evaluated based mainly on p4EBP1, and pS66 where a tumour is considered positive if certain percentage of cells is expressing the investigated marker. Similar analyses were performed for other compartments. B) Example images from the multiplexed dataset utilised in this study showing different markers from a selected tumour. Markers that showed no expression in the shown tumour are not shown.

Our primary focus was on the phosphorylation of S6 and 4EBP1, which serves as indicators of mTOR pathway activation (pS6 and p4EBP1 respectively). To systematically assess mTOR activity across different cell types in the tumour, we classified different cell types based on the epithelial marker pan-cytokeratin, stromal cell marker (smooth muscle actin (SMA)) and the immune markers (CD3, CD8, CD20, CD68, CD79) (Fig. 1A and Methods). Then we identified cells that are positive for either pS6 and p4EBP1 or both (Methods). Within the tumour, pS6 was expressed in 40% of epithelial cells, 20% of stromal cells and 40% of immune cells on average (Fig. 2A). Similarly, p4EBP1 is expressed in 40% of epithelial cells, 15% of stromal cells and 41% of immune cells per tumour on average (Fig. 2B). Notably, mTOR activity varies between tumours. While most tumours have pS6 positive cells in different tissue compartments, only 62%-70% of tumours have p4EBP1 positive cells (Fig. 2C-D). The lack of 4EBP1 phosphorylation in some tumours cannot be explained based on the level of pS6, S6, 4EBP1, or AKT as no correlation is observed (Fig. 2E-H). These results indicate that while mTOR is most active in epithelial cancer cells, it is also frequently active in immune and stromal cells and that p4EBP1 levels vary between tumours.

**Fig. 2.**
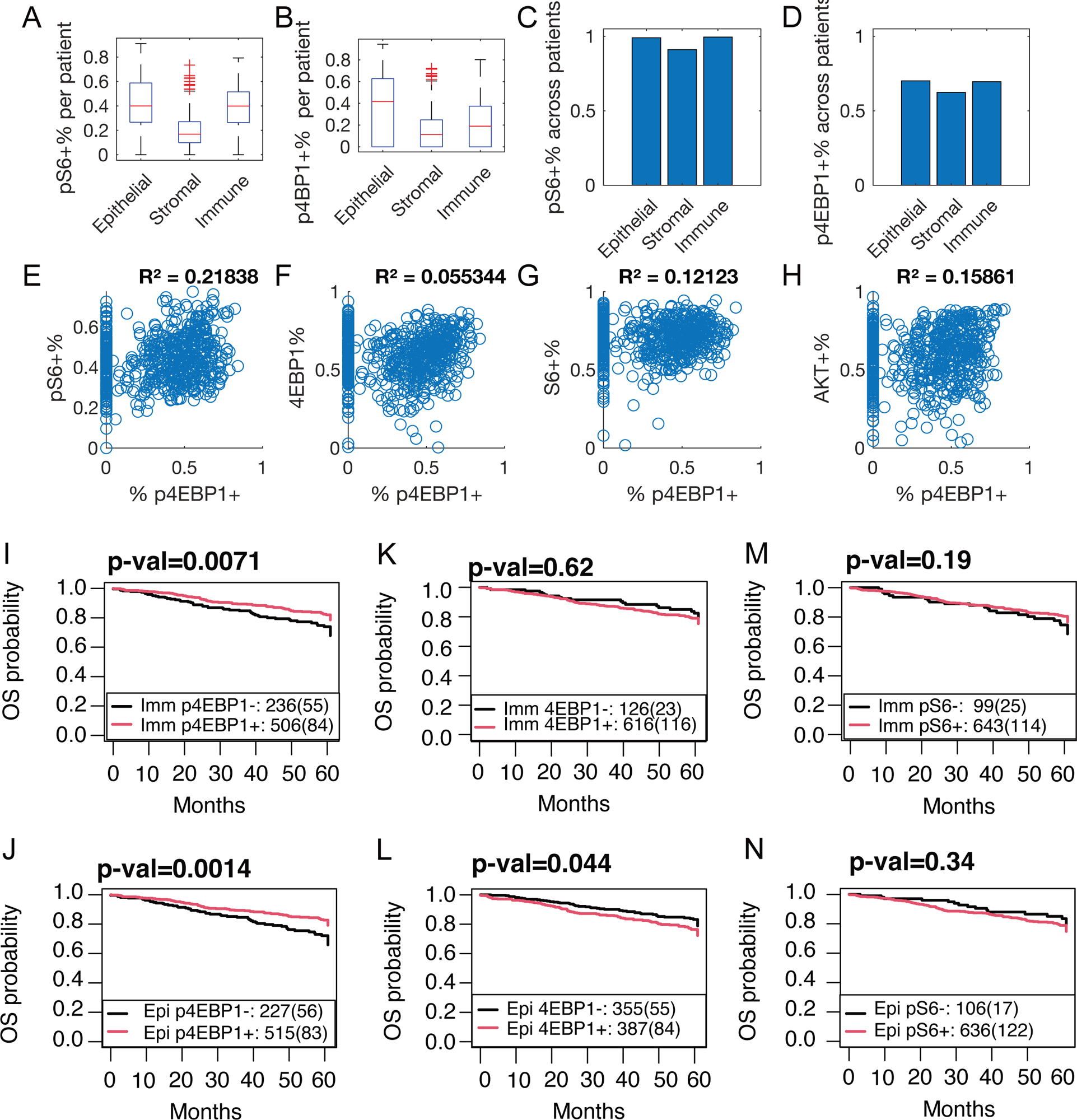
Analysis of mTOR activity in different tissue compartments and its association with survival. A-B) The distribution of the percentage of positive pS6 (A) and p4EBP1 (B) cells in different tumours. C-D) The proportion of tumours that are positive for pS6 (C) and p4EBP1 (D) where a tumour is considered positive for these markers if at least 5% cells were positive in that tumour. E-H) The relationship between the percentage of p4EBP1+ cells and pS6+ I, 4EBP1+ (F), S6+ (G), and AKT+ (H) where Pearson correlation is indicated on the top. I-M) Association between survival and immune signalling activity based on Kaplan Meier analysis. J-N) Association between survival and epithelial signalling activity based on Kaplan Meier analysis.

To determine the impact of mTOR activity in different cell types on patient outcomes, we performed Kaplan Meir survival analysis. We found that expression of p4EBP1 in immune (Imm^p4EBP1+^) or epithelial cells (Epi^p4EBP1+^) but not in fibroblasts is associated with better overall survival (Imm: pval=0.0071 and Epi: 0.0014, Fig. 2I-J and Supplementary Fig. 1A). The association between p4EBP1 and survival is consistent regardless of the cut-offs used to define p4EBP1+ tumours based on the percentage of p4EBP1+ positive cells (Methods and Supplementary Fig. 1B-C). Interestingly, 4EBP1 expression is associated with poorer outcomes only when expressed in the epithelial compartment (p-val = 0.044 Fig. 2K-L). We did not observe an association between pS6 expression and survival (Fig. 2M-N). As the phosphorylated form of 4EBP1 inhibits its action on EIF4, we investigated whether p4EBP1 association with survival is altered when combined with 4EBP1 expression levels. We observed that expression of 4EBP1+ in immune compartment does not affect the association between Imm^p4EBP1+^ tumours and survival (Supplementary Fig. 1D). On the other hand, the association between epithelial p4EBP1 and survival diminishes when 4EBP1 is also expressed in epithelial cells even at low levels (Supplementary Fig. 1E). These findings suggest that inhibition of 4EBP1 through its phosphorylation could provide prognostic advantage and its association with survival in epithelial cells is dependent on the levels of unphosphorylated 4EBP1.

We further examined the association between mTOR and various clinical variables. The number of Imm^p4EBP1+^ cells did not correlate with age or grade. However, we observed a significant association between Imm^p4EBP1+^ tumours and stage (chi-square test *p-val*= 5.8678e-09). Specifically, the number of Imm^p4EBP1+^ cells was highest in Stage 2 but lowest in Stage 3 (Supplementary Figure 1F). Cox multivariate regression analysis showed that the association between Imm^p4EBP1+^ and survival is independent from grade, gender, or age but not from tumour stage (Supplementary Table 2). Nonetheless, Wald test indicated that even though Imm^p4EBP1+^ is not independent predictor of tumour stage, these two variables collectively have highly significant effect on survival with high concordance (Wald test *p-val* = 1.2E-12, concordance =0.69).

### Characterisation of signalling and TME in Imm^p4EBP1+^ tumours

To identify signalling activities associated with Imm^p4EBP1+^ tumours, we investigated differentially expressed markers in tumours with p4EBP1+/- immune cells. The top 30 significantly different markers were selected (Methods). The levels of LaminA/C, FOXO3a, EZH2, EGFR, ALDH1, NaKaTPase, FOXO1, and p21 in immune cells are lower in Imm^p4EBP1+^ tumours compared to Imm^p4EBP1-^ tumours. However, the level of these proteins in cancer cells is comparable between Imm^p4EBP1+^ and Imm^p4EBP1-^ tumours (Fig. 3A). The cystine/glutamate antiporter xCT or SLC7A11 is significantly reduced in both immune cells and epithelial cells in Imm^p4EBP1+^ tumours. Additionally, Imm^p4EBP1+^ tumours exhibit a decrease in cMet, CyclinB1 and Glut1, beta-actin in both immune and epithelial compartments. Furthermore, positive tumours exhibit higher levels of ERK and downstream signalling of MAPK such as GSK3a, p38MAPK, pMAPAPK2, pS6235, pERK, NDRG1, MLH1, MSH2, p53, and GSKbeta in the epithelial compartment. A similar trend is observed when comparing p4EBP1+ and p4EBP-immune cells within the same tumour, except for FOXO1 and NaKaTPase. These results suggest that p4EBP1 activation is associated with lower proliferation, reduced metabolic activity, and decreased stemness markers but higher levels of DNA repair proteins.

**Fig. 3.**
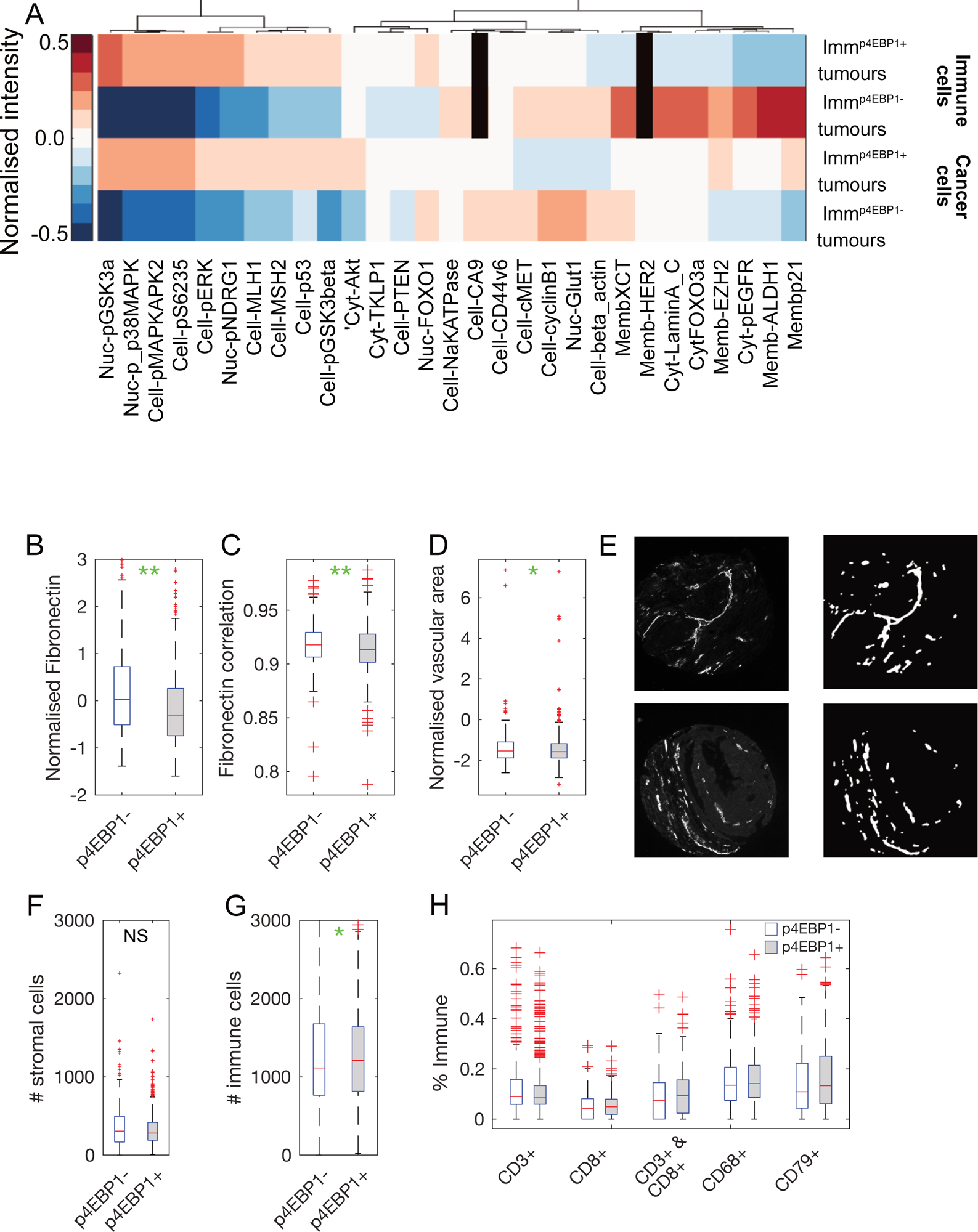
Cell signalling and TME properties in Imm^p4EBP1+^ tumours. A) Top differentially expressed proteins in immune and epithelial compartments in Imm^p4EBP1+^ tumours. Black rectangles indicate markers that are not known to be expressed in immune cells (Methods). (B) Fibronectin levels in Imm^p4EBP1-^ versus Imm^p4EBP1+^ tumours based on normalised intensity (p-val=1.4e-5). (C) Correlation between pixels in fibronectin images in Imm^p4EBP1-^ versus Imm^p4EBP1+^ tumours (p-val=0.0013). D) Difference in vascular area in Imm^p4EBP1-^ versus Imm^p4EBP1+^ tumours. E) Example images showing vessel segmentation for measuring vessel area based on CD31 marker. F) Number of stromal cells in Imm^p4EBP1-^ versus Imm^p4EBP1+^ tumours. G) Number of immune cells in Imm^p4EBP1-^ versus Imm^p4EBP1+^ tumours (p-val = 0.0083). H) Percentage of different immune cells types in Imm^p4EBP1-^ versus Imm^p4EBP1+^ tumours (* indicates p-val <0.05).

Next, we compared the TME between Imm^p4EBP1+^ and Imm^p4EBP1-^ tumours. We found that fibronectin and vascularisation are reduced in tumours with Imm^p4EBP1+^ (Fig. 3B-E and Methods). Importantly, these changes were not due to differences in the number of stromal cells (Fig. 3F). We also observed an increase in total number of immune cells including cells positive for both CD3 and CD8 (p-val<0.05 and Fig. 3G-H). These results suggest that p4EBP1 activity is associated with immune activation which may be sustained by lower fibronectin disposition.

### Association between Imm^p4EBP1+^ cells and tumour composition

Next, we sought to investigate whether Imm^p4EBP1+^ tumours are associated with certain signalling phenotypes in cancer cells. To account for all potential cellular interactions, we performed clustering of all tumour cells, totalling 1,818,510 cells, based on single-cell protein measurements. We identified 10 single-cell clusters (scClusters) using K-means (Fig. 4A-B and Methods). Kaplan Meier survival analysis revealed that scCluster 3, 4, 7, and 9 are associated with worse overall survival (OS) while scCluster 1 and 5 were associated with better OS and progression-free survival (PFS) (Fig. 4C and Supplementary Fig. 2C). Interestingly, even though scCluster 4 and 7 represents less than 5% of tumour cells within most tumours, they are associated with worse survival (Fig. 4B). scCluster 4 correlated most with scCluster 3 and 9 followed by 7 making it difficult to determine which of these clusters is the primary driver of cancer progression (Fig. Supplementary 2A). In general, clusters that are associated with worse survival are correlated negatively with immune infiltration and sometimes positively correlated with proportion of CD68 cells (Fig. 4F). Conversely, clusters that are associated with better survival, are positively correlated with the proportion of immune cells.

**Fig. 4.**
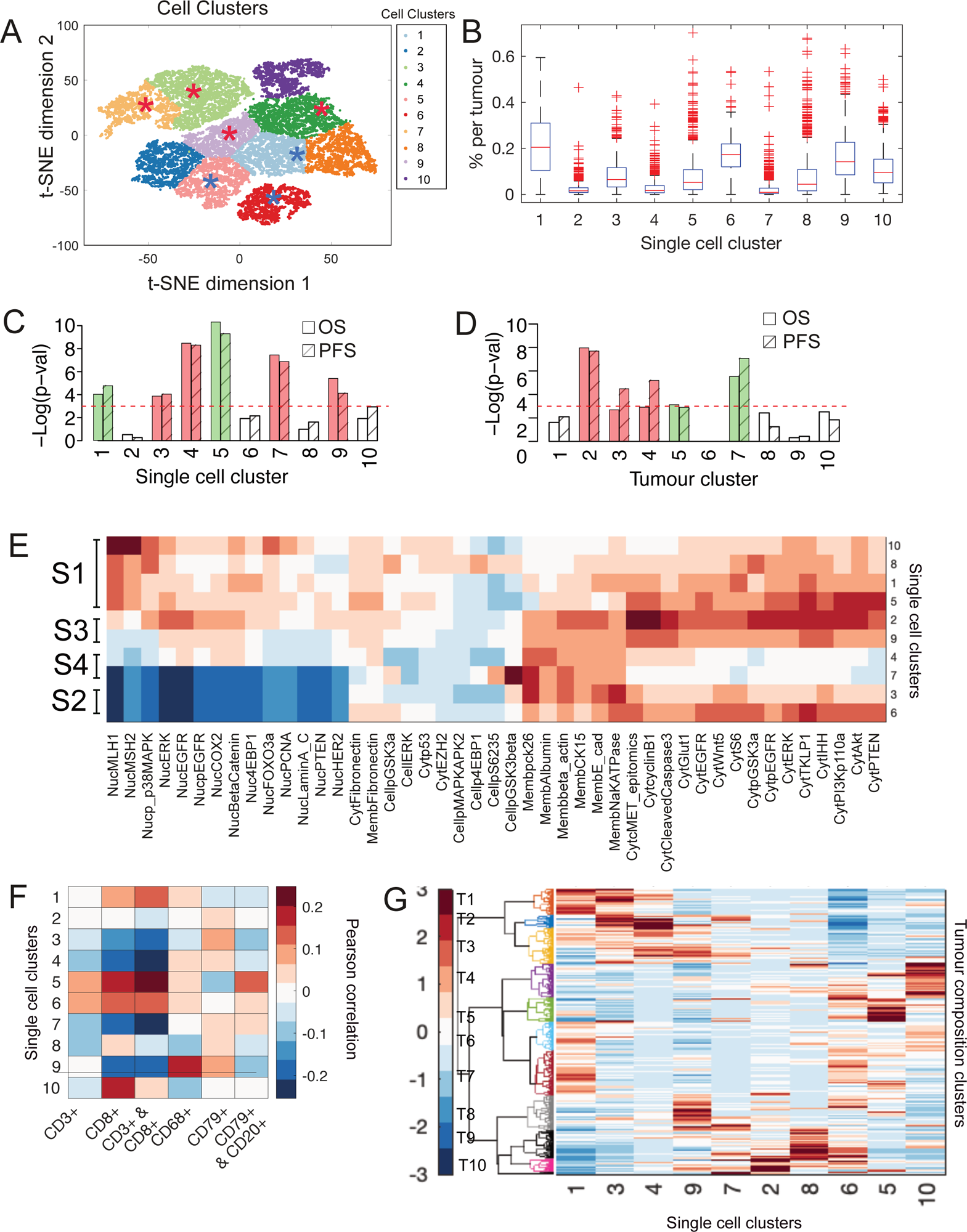
Association between Imm^p4EBP1+^ tumours and cancer cell signalling phenotypes. A) Visualisation of single cancer cells using t-SNE and their Clustering based on their signalling measurements using K-means. Red stars indicate the clusters associated with worse overall survival and blue stars indicate clusters associated with better overall survival. B) Proportion of various scClusters in different tumours. C) Association between enrichment for a certain scCluster and survival. Shown is -log of p-value based on Kaplan Meier survival analysis. The red line indicates p-value = 0.05 where higher values indicate lower p-values. Red bars indicate an association with worse survival and green bars indicate an association with better survival. D) Similar to C but for tumour clusters. E) Average signal per scCluster for most differentially expressed markers show 4 key signatures labelled by S1-S4. E) Pearson correlation between the percentage of each scCluster and the percentage of various immune cells in different tumours. G) Clustering of tumour heterogeneity profiles which represent the normalised proportion of different scClusters.

To determine key signalling state of each scCluster, we calculated the average signal per scCluster (Fig. 4E). This revealed four distinct signalling signatures. Signature 1 includes scCluster 10, 8, 1, and 5 and exhibits high levels of DNA repair proteins (MSH2, MLH1) and proliferation suppressors (phospho-p38 and nuclear FOXO3a). On the other hand, Signature 2 (scCluster 3, 6) is characterised by the lowest levels of DNA repair proteins (MSH2, MLH1, PCNA and Lamin A/C) and proliferation suppressors (Foxo3a, PTEN and p38). These clusters also exhibit a decrease in nuclear EGFR and ERK, pEGFR and HER2, suggesting altered transcriptional activities. Signature 3 (scCluster 2, 9) and Signature 4 (scCluster 4, 7) are similar to Signature 1 but express a lower level of MSH2 and MLH1. However, Signature 4 also exhibits the lowest levels of GLUT1, AKT, PI3Kp110, WNT, CyclinB1, cytoplasmic ERK and EGFR. scCluster 7 exclusively exhibit higher level of pGSKbeta and pS6. Only scCluster 8 show higher p4EBP1 activity in cancer cells compared to other clusters. The enrichment for Imm^p4EBP1+^ cells in various scClusters was measured using Jaccard index which confirmed that scCluster 8 is the most enriched with moderate Jaccard index of 32% followed by scCluster 2, 6, and 7 (Supplementary Figure 2). We did not observe that Imm^p4EBP1+^ cells were particularly depleted in presence of any scClusters. These results suggest that p4EBP1 activity in immune cells is not strongly associated with a specific signalling signature in cancer cells.

As most tumours are composed of a mixture of scClusters, we asked whether tumour composition, rather than the presence of specific scClusters, could distinguish tumours with Imm^p4EBP1+^ cells. To characterise tumour composition phenotypes, we clustered tumours based on the normalised percentage of each scCluster resulting in 10 tumour clusters (Fig. 4G and Methods). The association between tumour clusters and survival could be primarily explained by their single-cell composition. However, we also observed that, on some occasions, the occurrence of certain clusters had the converse effects on survival depending on the presence of other clusters. For example, colocalization of scCluster 6 (low DNA repair) and scCluster 10 (high DNA repair) in tumourCluster 4 associates with worse PFS while colocalisation of scCluster 6 and 5 in tumourCluster 5 associates with better PFS. Similarly, while tumourCluster 1-3 were all enriched for scCluster 3, only tumourCluster 2-3, which are also enriched for scCluster 4 were associated with worse survival. Again, we did not observe that Imm^p4EBP1+^ tumours were associated with a certain tumourCluster.

### Graph neural network for deciphering cellular networks responsible for Imm^p4EBP1+^ cells

The analysis of single cancer cells highlighted the need for a more comprehensive approach that account for different cell types and their signalling state. To this end, we utilised a graph neural network (GCN) in which we constructed a network of cells to represent the spatial relationship between cells in the tumour^21^. Specifically, we defined cells as nodes within a network, with edges representing interactions between neighbouring cells within a 50-pixel radius (Fig. 5A). Each cell node in the graph incorporated node features representing its signalling and morphology measurements. This network was then fed into a GCN to predict Imm^p4EBP1+^ versus Imm^p4EBP1-^ tumours (Fig. 5B and Methods). We achieved Area Under the ROC Curve (AUC) of 82%, an Average Precision (AP) of 91%, and an accuracy of 75% (Methods and Supplementary Fig. 3A). GCN also provided confidence scores for the predictions. Notably, GCN exhibited high confidence (absolute confidence score > 0.3) in 60.72% of the correct predictions (Fig. 5D-E). Conversely, for the incorrectly predicted tumour status, GCN showed significantly lower confidence, with only 14.6% of those tumours having an absolute confidence score greater than 0.3 (Fig. 5E). That GCN can predict mTOR status in immune cells, indicate a role for the TME in driving this phenotype.

**Fig. 5.**
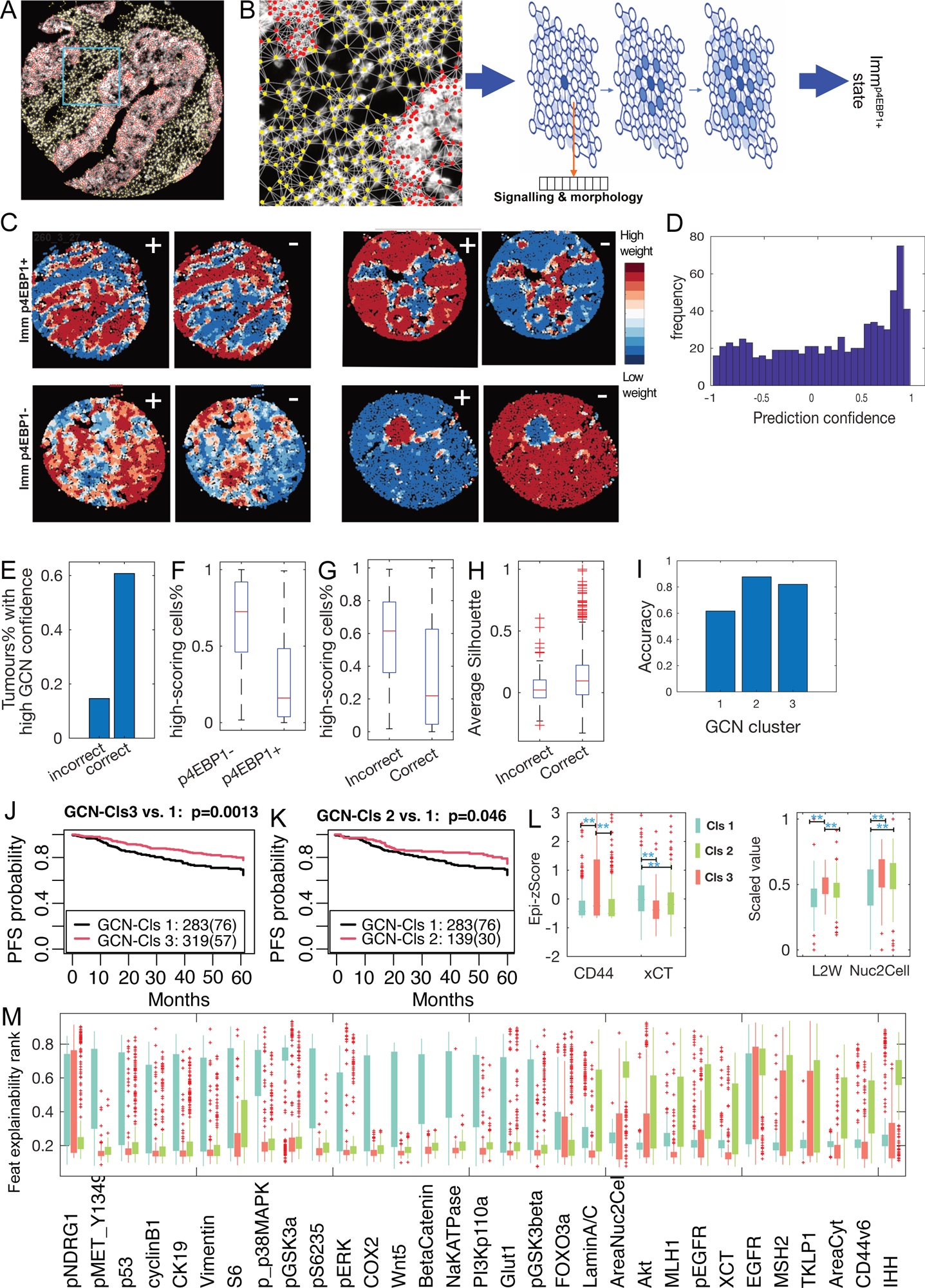
Prediction of Imm^p4EBP1+^ tumours and identification of relevant cellular subpopulation using graph neural networks. A) Cells projected as a network based on their spatial distance. B) Representation of GCN classifier through message passing between cells to predict p4EBP1 state. C) Visualisation of node/cell scores contributing to the positive class versus negative class for 4 tumours. Red indicates a positive contribution and blue indicates a negative contribution. D) Distribution of GCN prediction confidence. E) The percentage of tumours that were predicted correctly or incorrectly by GCN and achieved more than 0.3 confidence score. F-G) The percentage of high scoring nodes in Imm^p4EBP1+^ tumours versus Imm^p4EBP1-^ tumours (F), or correctly and incorrectly predicted tumour labels (G). H) Clustering of high-scoring nodes based on silhouette index in correctly and incorrectly predicted tumour labels. I) Accuracy of GCN across different GCN clusters. J) Association between PFS and GCN cluster 1 versus 3 using Kaplan Meier survival analysis. K) Association between PFS and GCN cluster 1 versus 2. L) Right: CD44 is significantly higher in GCN cluster 2 while xCT is significantly lower. Left: Cell elongation (Length/width ratio) and nuclear/cytoplasmic ratio is significantly higher in GCN cluster 2. ** indicates p-val<0.005. M) The distribution of feature ranks in the top selected features by GCN explainer across different GCN clusters. CD44 and cell morphology are among the most distinct features between GCN cluster 2 and 3.

Next, we sought to determine the extent to which Imm^p4EBP1+^ contributed to the predictions. GCN assigns two scores to each cell, representing their contribution to the negative and positive classes (i.e. Imm^p4EBP1-^ versus Imm^p4EBP1+^). Visualising node weights for the negative and positive classes revealed that they tend to be inversely correlated (Fig. 5C and Methods). We also notice that sometimes a gradient of weights was learnt while in other cases homogenous weights were learnt for the contributing cells. Cells with high weights in the positive class were defined as high-scoring cells. Interestingly, we found that 75.05% of immune active mTOR cells were assigned a high weight by GCN. These cells represent only 8.95% of high-scoring cells whereas other high-scoring cells constituted 26.13% immune, 7.4% stromal and 66.47% epithelial cells. This provided confidence that the network can detect the relevant signature despite that p4EBP1 measurements not being included in the cells features and GCN was only given the core-level labels.

To better understand GCN operations, we analysed the distribution of high-scoring cells. We observe that tumours varied in the number and localisation of high-scoring cells contributing to the positive class. In some tumours, only a small number of cells have high weights while in other tumours, the majority of cells contributed to the prediction (Fig. 5C). Interestingly, the number of high-scoring cells contributing to Imm^p4EBP1+^ label was significantly less in the Imm^p4EBP1+^ tumours compared the Imm^p4EBP1-^ tumours (Fig. 5F). Similarly, the number of high-scoring cells was significantly less when GCN correctly predicted Imm^p4EBP1+^ class (Fig 5G). Top-scoring cells tend to be clustered together based on the silhouette index which was also higher when the network made correct predictions (Fig. 5H). One possible explanation for that is only a subset of clustered cells was relevant for the positive class, but the network tends to consider the state of most cells when predicting the negative class.

We utilised GCN explainer to identify the features that contributed most to p4EBP1 activation in immune cells in different tumours^22^. The algorithm allowed us to weigh the contribution of each feature to the prediction of each tumour (Methods). We identified the top-ranking features across all tumours (scored >0.4 in at least 200 tumours). This resulted in 81 features that were broadly similar to what we identified by the differential analysis in Figure 3A (Fig. 5M). Clustering of tumours based on GCN feature rankings identified that tumours with and without Imm^p4EBP1+^ cells can be distinguished based on different set features (Supplementary Fig. 3B). Interestingly, this clustering revealed that tumours with Imm^p4EBP1+^ cells can be divided into two subgroups (GCN-Cls2 and GCN-Cls3) (Supplementary Fig. 3B). These two clusters were generally similar in terms of number of Imm^p4EBP1+^ cells, accuracy, number of Imm^p4EBP1+^ cells, confidence, and the number of high-scoring cells (Fig. 5I and Supplementary Fig. 3E-G). Interestingly, GCN-Cls3 have a more significant association with better PFS compared to GCN-Cls2 (*p-val*= 0.0015 and 0.045 respectively and Fig. 5J-K) but only GCN-Cls3 was highly associated with better OS (*p-val*= 0.0007 and Supplementary Fig. 3D). This is more significant when compared to p-value of 0.007 when both GCN-Cls2 and 3 were combined. Importantly, GCN-Cls3 and 2 are not associated with tumour stage or grade (Supplementary Fig. 1H). Therefore, GCN allows further stratification of tumours with Imm^p4EBP1+^ cells based on holistic analysis of spatial protein activity.

To further understand the association between p4EBP1 activity in immune cells and patient outcome, we compared the values of the top predictive features across GCN-clusters based on GCN explainer (Fig. 5M). In GCN-Cls1, these included phospho-p38, pMET, PI3Kp100, and the adhesion protein BetaCatenin, NaKaTPase (Supplementary Table 3). Only a few features have high ranking for GCN-Cls2 and these overlapped with GCN-Cls3. In GCN-Cls3, predictive markers include xCT, MSH2, MLH1, pEGFR, EGFR, and AKT which is consistent with our differential analysis (Fig. 3B). Interestingly, markers of stemness such as CD44, IHH and Lamin A/C, as well as morphological features, such as the ratio of nuclear/cell area exhibited the highest predictiveness for GCN-Cls3, which was not observed in the case of GCN-Cls2 (Fig. 5M). The values of these variables indicate that GCN-Cls3 has a more differentiated morphology and lower CD44 than GCN-Cls2 (Fig. 5L, Supplementary Fig 3C, and p-val<0.005). Based on these results we propose that Imm^p4EBP1+^ offer more significant benefits to the patients when the cells in the tumour maintain a well differentiated state, and their positive impact diminishes as epithelial cells acquire a more stem-like state.

### Identification of molecular mechanisms using spatial transcriptomics

To elucidate the mechanism and functional relevance of p4EBP1 activation in immune cells, we conducted spatial transcriptomics using the GeoMx Digital Spatial Profiler from NanoString on an independent cohort of patients (Fig. 6A-D and Methods). We were able to harvest RNA from immune cells in 20 tumours that showed various levels of p4EBP1 activity in immune cells (4 negative, 4 low, 8 medium, and 4 high). Additionally, we harvested RNA from PCK+ tumour cells as a control. We confirmed that GeoMx approach successfully separated transcripts from these distinct populations with immune segments expressing CD45 and lacking expression of PCK, while the opposite is true for PCK+ segments (Fig. 6E-G).

**Fig. 6.**
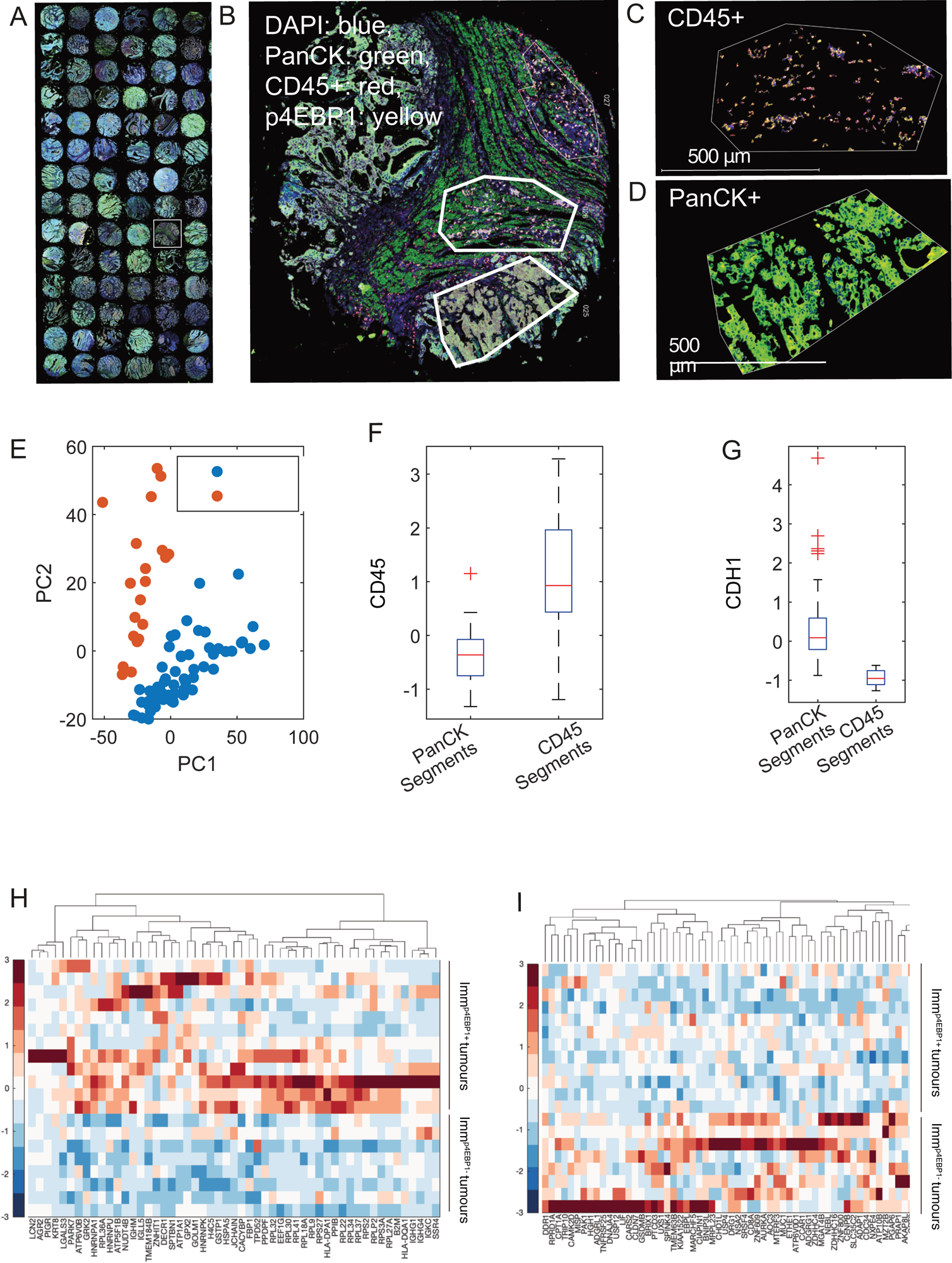
Determination of the functional relevance of p4EBP1 activity on immune function and tumour microenvironment. A) TMA stained for DAPI (blue), PCK (green), p4EBP1 (yellow), and CD45 (red). B) Inset of the region highlighted in A. C-D) Polygons are manually drawn to select immune cells (CD45+) versus tumour cells (PCK+). E) RNA sequencing data of profiled regions are visualised in PCA space and show that CD45+ regions are clearly separate from PCK+ regions. F-) CD45 and PCK expression in CD45+ regions versus PCK+ regions respectively. H) Upregulated genes in immune regions in Immp4EBP1+ tumours. I) Downregulated genes in immune regions in Imm^p4EBP1+^ tumours.

Differential expression analysis revealed variations between tumours with and without Imm^p4EBP1+^ cells. Imm^p4EBP1+^ tumours exhibited upregulation of genes involved in transcription including ribonucleoprotein (16 genes), translation (13 genes), and RNA binding which is expected given the role of 4EBP1 in transcription regulation (Fig. 6H and Table EV4). There was also a higher expression of genes associated with immune activation such as those involved in immunoglobulin/major histocompatibility including IGHM, IGLL5, HLA-DQA1, and HLA-DPA1, and lymphoblast (15 genes) (*p-val* < 0.05). Similar enrichment results were obtained when considering only the tumours negative for p4EPB1 versus other tumours (results not shown). On the other hand, downregulation was observed in genes associated with proliferation genes, such as AURKA, DDR1 and MISP as well as collagen receptors, such as ADGRG1, DDR1 and other adhesion proteins such as MUC1 (Fig. 6I). This suggests that p4EBP1 activity in immune cells is associated with activation of immune cells and suppression of proliferation and ECM interactions.

Interestingly, we found that these genes are specifically differentially expressed in the immune compartment and not in the epithelial compartment. The small number of differentially expressed genes in the tumour region did not show clear significant enrichment. However, we do observe that TGFBI, and the laminin subunit 2 (LAMC2) are reduced in the epithelial region in Imm^p4EBP1+^ tumours. Given the low number of differentially expressed genes in the tumour region when p4EBP1 is high or low in the immune compartment, we speculate that Imm^p4EBP1+^ is not primarily driven by certain properties in the tumour.

## Discussion

The tumour microenvironment presents numerous interactions with the tumour, affecting its progression and evolution. Typically, signalling in the tumour and the surrounding environment is often studied separately in the different compartments. Single-cell and spatial biology techniques provide enabling technologies for studying complex and heterocellular signalling in the TME. In particular, multiplexed imaging offers the unique advantage of measuring spatial protein activities including phosphorylation events, at the single cell level. Here we show how highly multiplexed imaging can advance our understanding of complex mTOR cell signalling.

Several approaches have been proposed for analysing multiplexed datasets and understanding the association between tumour heterogeneity and microenvironment signatures with patient prognosis^3,4,23,24^. Unlike unsupervised approaches for detecting cellular communities and local neighbourhood^22,23^, graph neural networks allow the consideration of larger spatial regions through iterative message passing within the tumour network. They also allow the identification of relevant features given the output class which is in our case the mTOR signalling state. Our analyses show that GCNs complement other single cell and heterogeneity analyses, enabling the selection of relevant cellular subpopulations and patients stratification.

Our findings indicate that a more favourable patient prognosis in Imm^p4EBP1+^ tumours is influenced by a combination of various factors within the TME, including reduced fibronectin deposition, decreased vascularization, and a higher number of immune cells. Spatial transcriptomics data further confirms that p4EBP1 activity in immune cells is associated with immune activation. We also found that Imm^p4EBP1-^ tumours exhibit higher levels of CD44 leading which could lead to suppression of immune cells by through its interaction with TGF-beta that simulate ECM production.

GCN analysis further supported the link between stemness and 4EBP1 phosphorylation in immune cells. Specifically, by integrating the influence of tissue morphology, cellular communication, and signalling states in different tissue compartments, GCN identified two sub-groups of Imm^p4EBP1+^ tumours. GCN-Cls3 with more differentiated morphology and GCN-Cls2 with more stem-like properties. This division achieves a better stratification of patient outcomes where GCN-Cls3 has a more significant association with better OS and PFS. These results suggest stem programs, such as EMT, antagonise the anti-tumour role of Imm^p4EBP1+^ and gradually lead to further progression of tumours and inhibition of 4EBP1 phosphorylation.

While our study focused on p4EBP1 expression in immune cells, our results also imply that p4EBP1 activity in epithelial cells could also have a positive effect on patient prognosis depending on 4EBP1 expression. Such positive effect could be attributed to the inhibition of 4EBP1 through its phosphorylation as the expression of 4EBP1 in cancer cells counteracted the impact of p4EBP1 effect on survival. This finding could provide explanation of the conflicting studies on the effects of p4EBP1 on patient survival.

Spatial transcriptomics suggest that the key transcriptional differences in Imm^p4EBP1+^ and Imm^p4EBP1-^ tumours are mainly in the immune compartment with no clear signature in the epithelial compartment except to a few genes. This suggests that the clinical benefit of p4EBP1 expression in immune cells is driven by immune response rather than signalling in the epithelium. However, stemness programmes could impact p4EBP1 indirectly through ECM remodelling.

In summary, we propose that mTOR plays a dual role, supporting cancer growth and proliferation on one hand while exerting anti-tumour activity when active in immune cells on the other hand. Although, many studies commented on these different aspects in diseased and healthy cells, our work is the first integrated evidence to support the different roles for mTOR in the TME. Our study offers a potential explanation of the limited progress of mTOR inhibitors and therefore could have a significant implication for the use of mTOR inhibitors in therapy. Most importantly, our study provides a framework for studying complex cell signalling and accounting for cell interactions within different TME components using multiplexed imaging and advanced analysis methods.

## Methods

### Multiplexed Dataset

Multiplexed immunofluorescence staining of the colon cancer TMAs was performed as previously described^1^ using Cell DIVE™ technology (Leica Microsystems, Issaquah, WA). After de-paraffinization and a two-step antigen retrieval, the FFPE slides were stained with DAPI and imaged in all channels of interest to acquire background autofluorescence (AF) of the tissue. This was followed by indirect detection and/or direct conjugate antibody staining of up to 3 markers per round plus DAPI, dye deactivation, and repeat staining to collect images of all planned biomarkers. Multiplexed images are automatically registered and processed for illumination correction and autofluorescence subtraction during each round of imaging using the Cell DIVE image processing workflow. (Supplementary Table 1) as previously reported in Gerdes et al., and Uttam et al^1,3^.

### Image pre-processing and analysis

Our analysis builds on custom image processing and segmentation algorithms from Gerdes *et al.,*^25^. In the present study, we identified different types of immune cells and maker expression were determined based on an adaptive thresholding method with results qualitatively validated. Intensity thresholds were adjusted when a high level of noise was detected. A similar approach was applied to determine cells that are positive for mTOR markers. Stromal and epithelial regions were defined based on pan-cadherin (PCK), E-cadherin and SMA expression, utilising a combination of adaptive thresholding and dilation/erosion image operations. Each cell was then assigned either to stromal or epithelial region as well.

### Fibronectin and vessel analysis

Adaptive thresholding was applied to CD31 images to delineate vessel regions, which were subsequently analysed to obtain vessel area. This was normalised to stromal area within the tumour. To quantify the structure fibronectin, we measured the correlation between different neighbouring pixels as a proxy for fibronectin structure.

### mTOR status

A tumour is defined to be Imm^p4EBP1+^ if at least 5% of immune cells in the tumour express p4EBP1. Based on this cut-off, ∼70% of tumours were considered positive for Imm^p4EBP1+^ (Fig. 2D). Such a threshold ensures robustness to staining artifacts. We confirmed that the association between Imm^p4EBP1+^ and survival is not sensitive to a certain threshold, by testing several thresholds. We found that Imm^p4EBP1+^ is consistently associated with survival probability based on Kaplan Meier analysis.

### Data normalisation

In addition to autofluorescence subtraction across different rounds, we normalised intensity features by calculating their z-score per slide to eliminate batch effect.

### Differential protein expression analysis

To identify signalling differences between Imm^p4EBP1+^ and Imm^p4EBP1-^ tumours, we performed Kolmogorov-Smirnov (KS) non-parametric test. The 30 most significantly different features in the immune compartment and epithelial compartment were considered which included p4EBP1 and therefore was excluded. The relevant cellular compartment was chosen per protein and the difference in average marker intensity was visualised.

Markers that are not known to be expressed in immune cells were determined based on one or more of the following: 1) Qualitative assessment of protein expression in immune cell in colorectal cancer samples in Human Protein Atlas^26^, 2) Expression of the corresponding gene in immune cells based on immune proteome atlas^26^, or 3) literature documenting a role for the protein in immune function.

### Single-cell visualisation

Principal Component Analysis (PCA) was applied to normalised intensity features of single cells. Then the first 5 Principal Components (PCs) were further reduced in two dimensions using tSNE.

### Single cancer cell clustering

Normalised single cell measurements were clustered using k-means because of its scalability to a large number of cells. The algorithm was run with 10 different initialisations for 200 iterations using squared Euclidean distance. The initialisation with the best performance is chosen.

### Survival analysis for scClusters

A tumour is considered to be enriched for a certain scCLuster if the proportion of that cluster is greater than its 75^th^ quantile across all tumours.

### Tumour composition/heterogeneity clustering

To identify tumour composed of similar scClusters composition, we calculated the percentage of each scCluster in each tumour^27^. To ensure that scClusters with low frequency are not overlooked due to small distance difference (i.e. are rare across all the tumours), we z-scored the frequency of each cluster across all tumours. This results in heterogeneity profiles that represent the enrichment of each scCluster across all tumours (Fig. 4G). These profiles were then clustered using hierarchical clustering with ward linkage and Euclidean distance.

### Graph Convolutional Network (GCN) training

GCN is a type of neural network designed to work with graph-structured data, enabling it to account for spatial cell interactions. The input of GCN is a network of cells in a tumour along with cell futures. These networks were defined by representing each tumour as a graph where each cell is a node. Cell nodes were connected by edges if they are within 50 pixels within each other. Signalling and morphology measurements excluding p4EBP1 measurements were embedded into each node. The network is trained to predict two classes: 1) Imm^p4EBP1+^ tumours and 2) Imm^p4EBP1-^ tumours based only on single cell features and their spatial localisation. The network produces the probability of a tumour belonging to each class. Our approach also allows obtaining the weights assigned by the network to each cell which provide indication of the node importance to the output class. The network was trained using 5-fold cross validation (n=746).

### GCN-Clustering

GNNExplainer was run for 200 epochs and four hops to rank the most predictive node features for each tumour in regard to the output class. Tumours were clustered using hierarchical clustering with Euclidean distance and Ward linkage based on the z-scored ranks of node features. This clustering revealed three key clusters dubbed GCN-Cls1-3.

### Spatial Transcriptomics

Spatial transcriptomics were run on a tissue microarray of 125 cores and core size of 1.5 mm. The cells were stained for DAPI, pan-cytokeratin, CD45 and p4EBP1. Due to the limitation of GeoMx platform, only best cores were selected manually to harvest RNA from cancer versus immune compartment based on CD45 and PCK expression. 20 cores were identified to have a sufficient number of immune and epithelial cells (at least 50 cells) per DSP guidelines. These were considered for further analysis.

### Digital Spatial Profiling (DSP)

Spatial transcriptomic profiling was conducted on the NanoString GeoMx® platform according to manufacturer’s guidelines and recommendations. Tissue was manually processed with 20 minutes heat-induced epitope retrieval at pH 9 followed by 15 minutes of proteinase K (1μg/ml; Life Technologies) digestion at 37°C. RNA was hybridised in situ with the GeoMx® Human Whole Transcriptome Atlas Probe Mix. For visualisation, tissues were stained using immunofluorescent primary antibodies against PanCK (clone: AE1+AE3, conjugate: AF532; NanoString) and CD45 (clone: PD7/26+2B11, conjugate: AF594; NanoString), as well as unconjugated anti-p4EBP1 Thr37/46 (clone: 236B4; CellSignalling) and anti-rabbit secondary antibody (clone: ab150083; conjugate: AF647; Abcam). Nuclear staining was performed using SYTO13 (NanoString). Library preparation from the collected oligonucleotides was carried out according to manufacturer’s instructions and the sequenced on paired-end 150 NovaSeq600 with sequencing depth of 1200 million reads. Sequencing reads were aligned to the whole genome using DSP Analysis Suite and low quality reads were eliminated. Q3-normalised expression data was exported from the DSP Analysis Suite and analysed using MatLab.

### Spatial Transcriptomics analysis

We performed differential expression analysis using rnaseqdiff^28^. We identified genes that have the highest log fold change at cut-off of 0.75 in tumours with no or small number of p4EBP1+ cells in immune compartment versus medium to high number of p4EBP1+ cells in immune compartment.

### Author contribution

HS conceived the study with FG. HS designed computational and validation experiments, performed the data analysis, and wrote the paper. RZ and ME contributed to the analysis. AC, MJ, JH, and FI ran the spatial transcriptomics. All authors reviewed the paper.

## Acknowledgements

This work was funded by Sir Henry Wellcome Fellowship (Grant Number 204724/Z/16/Z) and Corpus Christi Small Grant Research Fund (University of Oxford, 2022). FG was supported by the National Cancer Institute of the National Institutes of Health under award number R01CA208179.

## Supplementary Figure Captions

**Supplementary Fig. 1.** Association between mTOR and survival. A) p4EBP1, pS6 and AKT signalling in stromal cells is not predictive of survival. B) p4EBP1 expression is significantly associated with overall survival (OS) even when expressed in different percentages of immune cells in the stroma confirming the robustness of this association. C) Same as (B) but in epithelial cells. D) When at least 5% of cells express p4EBP1, 4EBP1 expression in different percentages of immune cells generally does not impact the association between immune p4EBP1 and survival. E) The association between epithelial p4EBP1 and survival is dependent on the lack of 4EBP1 as 4EBP1 expression even in a small number of cells will diminish the correlation. Shown is -log of p-value based on Kaplan Meier analysis of OS. The red line indicates p-value = 0.05 where higher values indicate lower p-values. F) Percentage of Imm^p4EBP1+^ and Imm^p4EBP1-^ tumours across different tumour stages, grades and based on low and high age (68 was chosen as 50^th^ quantile).

**Supplementary Fig. 2.** A) Correlation between the percentage of various single cell clusters. B) Jaccard index of Imm^p4EBP1+^ tumours in the identified single cell clusters shows that these tumours can have variable signalling signature. C) Association between OS and scClusters. D) Association between OS/PFS and tumour clusters. Only clusters with significant associations are shown.

**Supplementary Fig. 3.** A) AUC and AP curves for GCN model. B) Clustering of features based on their GCN ranks reveals that GCN predict Imm^p4EBP1+^ tumours based on a different set of features which distinguish three tumour groups named GCN-clusters. Imm^p4EBP1+^ tumours can be divided into groups GCN-Cls 2 and 3. C) Z-score of CD44 and xCT levels in different GCN clusters in the immune compartment. ** indicates p-val<0.005. D) OS is significantly higher GCN-Cls3 versus GCN-Cls1. E) GCN-Cls 2 and 3 have a comparable number of Imm^p4EBP1+^ cells when considering correctly predicted tumours. F) GCN-Cls 2 and 3 have a comparable confidence prediction when considering correctly predicted tumours. G) GCN-Cls 2 and 3 have a comparable number of high-scoring cells, the average distance between high-scoring cells and their clustering based on silhouette index in correctly predicted tumours. H) Association between GCN-Cls2 and 3 and tumour stage and grade are not significant (chi-square test p-val=>0.09).

## Supplementary Tables

**Supplementary Table 1.** List of markers included in this study. More details are available in the original study^1^.

**Supplementary Table 2.** Cox Multivariate regression results.

**Supplementary Table 3.** List of top-ranked features by GCN explainers along the GCN cluster.

**Supplementary Table 4.** Enrichment of the differentially expressed genes in Imm^p4EBP1+^ and Imm^p4EBP1-^ immune tumours based on DAVID database.

