## Supplementary figures and images for "Decoding mTOR signalling heterogeneity in the tumour microenvironment using multiplexed imaging and graph convolutional networks"

### Supplementary Figures

#### Supplementary Figure 1

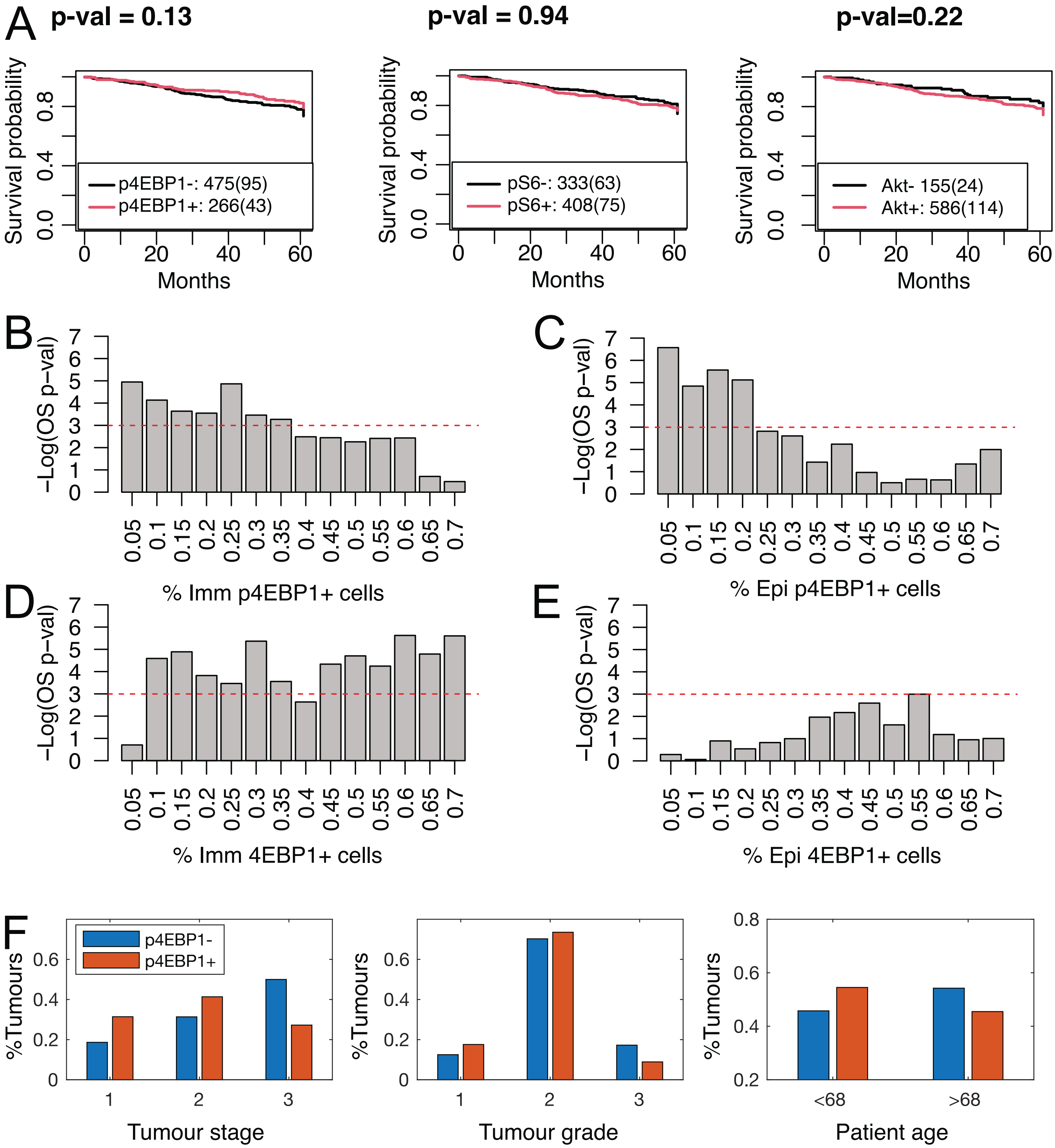

#### Supplementary Figure 2

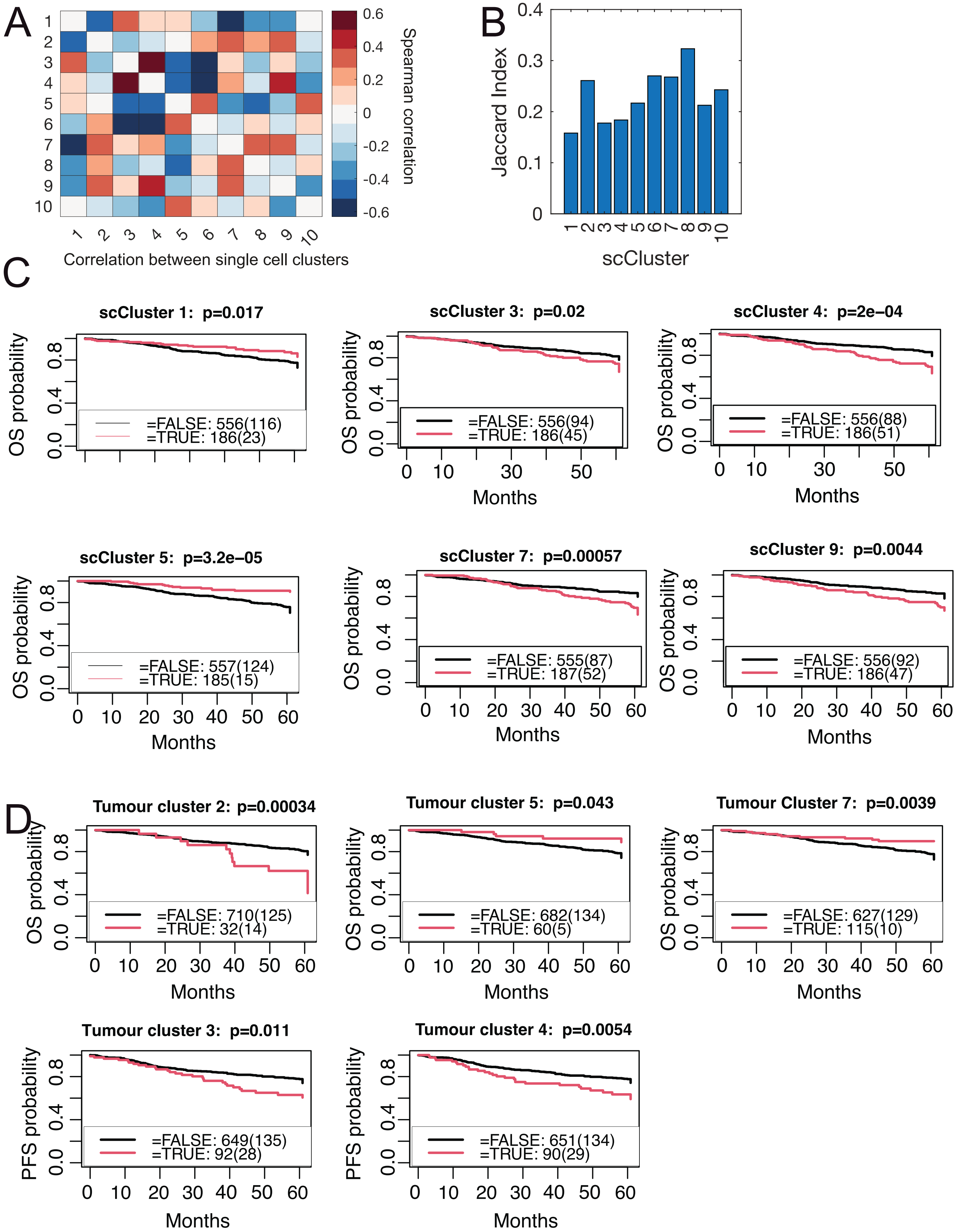

#### Supplementary Figure 3

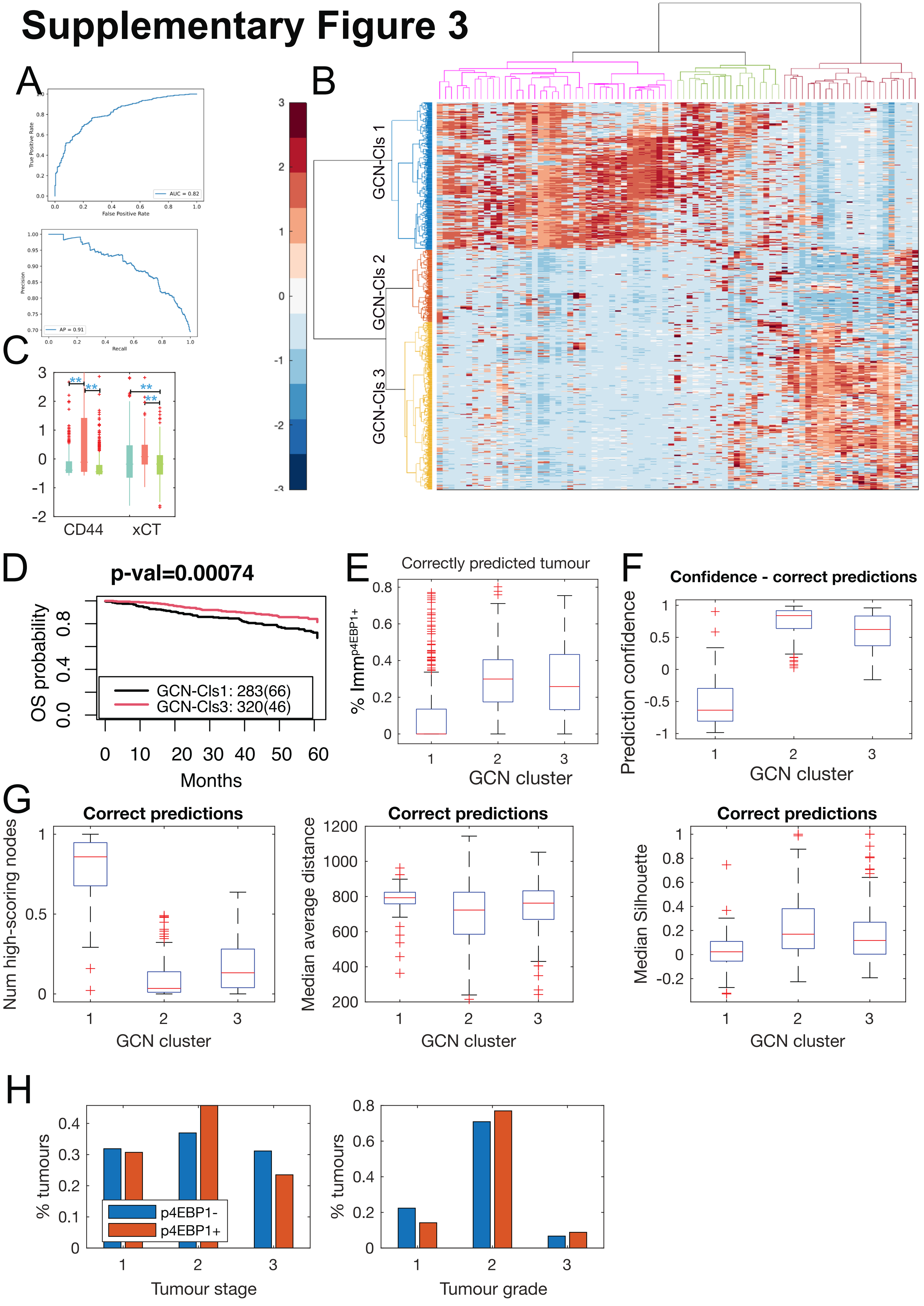
